# Calibration allows accurate estimation of the axonal volume fraction with diffusion MRI

**DOI:** 10.1101/2022.04.12.488014

**Authors:** Sebastian Papazoglou, Mohammad Ashtarayeh, Jan Malte Oeschger, Martina F. Callaghan, Mark D. Does, Siawoosh Mohammadi

**Affiliations:** Department of Systems Neuroscience, University Medical Center Hamburg-Eppendorf, Hamburg, Germany; Wellcome Centre for Human Neuroimaging, UCL Queen Square Institute of Neurology, University College London, UK; Department of Biomedical Engineering, Vanderbilt University, Nashville, TN, US; Institute of Imaging Science, Vanderbilt University Medical Center, Nashville, TN, US; Department of Radiology and Radiological Sciences, Vanderbilt University Medical Center, Nashville, TN, US; Department of Electrical Engineering, Vanderbilt University, Nashville, TN, US; Department of Neurophysics, Max Planck Institute for Human Cognitive and Brain Sciences, Leipzig, Germany

**Keywords:** Diffusion-weighted imaging, biophysical model, axonal water fraction, axonal volume fraction, unmyelinated axons, calibration, histology gold standard, g-ratio

## Abstract

Biophysical models enable the non-invasive estimation of microstructural tissue features of the central nervous system such as the axonal volume fraction using diffusion weighted imaging (DWI)-data. However, these models trade accuracy with complexity and demands for time-efficient data acquisition. In this study, we hypothesise that their accuracy can be improved substantially through biophysically motivated, linear calibration. We test this hypothesis in the context of axonal volume fraction estimation in four different DWI-models of different complexity using multi-modal data including ex-vivo diffusion MRI- and electron microscopy (EM)-data in mice with broad dynamic range, whereby the latter served as gold standard. We found that two calibration parameters, an offset accounting for the fraction of unmyelinated axons in severely hypomyelinated mice and a scaling accounting for the compartment-specific relaxation, substantially improved the accuracy of axonal volume fraction. Furthermore, we theoretically predict the scaling parameter, and demonstrate that similar accuracy improvement can be achieved for a subset of DWI-models, if the scaling parameter is fixed to the predicted value instead of estimating it on basis of data. This is of practical relevance because it allows to estimate the remaining offset calibration parameter from a limited amount of multi-modal data and thus makes the proposed method usable in human brain data.

## 1. Introduction

### All models are wrong but some are useful

This often cited aphorism attributed to the statistician George Box describes the dilemma that neuroscientists encounter when using biophysical models of the MR-signal as a non-invasive tool to infer microstructural tissue features. The most prominent model based on diffusion-weighted-imaging is Neurite Orientation Dispersion and Density Imaging (NODDI) which is often used to estimate the axonal volume fraction (*f*_*A*_). Accurate estimation of *f*_*A*_ of white matter (WM) is of great interest because it is required, e.g., for the computation of the g-ratio which is indicative of neuronal conduction velocity and thus of the functional integrity of WM fibres (Stikov et al., 2015; Campbell et al., 2018; Mohammadi & Callaghan, 2021). In vivo, MR-based *f*_*A*_ estimation is usually achieved by rescaling the axonal water fraction metrics from DWI-models with myelin volume fraction estimates from other non-DWI MR-techniques such as, e.g. proton density mapping or magnetisation transfer imaging (Stikov et al., 2015). The employment of non-DWI MR-techniques for estimating the myelin volume fraction is necessitated by the short transverse relaxation time of the macromolecular water, which renders it practically invisible to DWI.

A whole range of biophysical multi-compartment DWI-models, connecting the diffusion weighted signal with the axonal volume fraction, have been proposed such as, e.g., White Matter Tract Integrity (WMTI) (Fieremans et al., 2011), NODDI (Zhang et al., 2012), a reduced version of NODDI (NODDI-DTI) (Edwards et al., 2017), and most recently a model by Jespersen et al. (JESP) (Jespersen et al., 2018), which represents an extension of the WMTI-model. Each of these DWI-models is based on certain simplifying assumptions regarding white matter microstructure and the diffusion of water inside it, which result in different numbers of free model parameters. The JESP model has five free parameters, one intra-cellular (i.e. axonal) diffusivity parallel to the WM fibres, two extra-cellular diffusivities one parallel and one perpendicular to the fibres, the dispersion of fibres, and the axonal water fraction. Furthermore, due to the degeneracy of its solution, the JESP model possesses two solutions JESP+ and JESP-. WMTI assumes no fibre dispersion, i.e. WM fibres are assumed to be parallel inside a given voxel, which reduces the number of free parameters to four. NODDI is the only three compartment (four compartment ex-vivo) model and includes an isotropic, cerebrospinal fluid compartment in addition to the extra-cellular and the axonal compartments shared by all DWI-models of this study. The free parameters of NODDI are fibre dispersion, axonal water fraction and isotropic water fraction and, for the ex-vivo NODDI model, the signal fraction of the dot compartment (restricted water pool (Alexander et al., 2010; Panagiotaki et al., 2012)) for the ex-vivo NODDI model. The compartmental diffusivities are fixed and not independent to avoid overfitting. Finally, NODDI-DTI is the simplest model because it considers only WM voxels, in which the isotropic and the dot compartmment can be ignored. Therefore, it has only two free parameters, fibre dispersion and axonal water fraction. The remaining compartmental diffusivities are fixed as for NODDI.

Although the limitations of these biophysical DWI-models are well-known, they are increasingly used in clinical research and neuroscience, e.g. NODDI (Cox et al., 2016; Elliott et al., 2018; Genç et al., 2018; Donat et al., 2021) or WMTI (Coutu et al., 2014; Karolis et al., 2019). One important limitation is the fact that these models neglect different compartmental *T*_2_ values in the intra- and extracellular signal (Veraart et al., 2018; Lampinen et al., 2019; Gong et al., 2020; Frigo et al., 2021). Another limitation is that their sensitivity to the fraction of unmyelinated axons is unknown. While additional linear calibration has often been used to improve the accuracy of MR-based myelin metrics (Campbell et al., 2018; Jung et al., 2018; West et al., 2018b), the question about whether additional calibration can also improve the accuracy of DWI-based axonal water estimates has remained unanswered (Mohammadi & Callaghan, 2021). The signals of the aforementioned multi-compartment DWI-models are essentially modeled as the sum of signal contributions from the individual compartments (e.g. axonal, extracellular, isotropic compartments). The fraction of the signal of the axonal compartment is then usually taken directly as the metric for the axonal water fraction (*f*_*A*_). However, signal fraction and axonal water fraction are not truly interchangeable, because of the different transverse relaxation times (*T*_2_) in the compartments. Instead, the signal fractions are potentially weighted fractions whose weights depend on the compartmental *R*_2_-differences (*R*_2_ = 1*/T*_2_) and on the employed echo time *T*_*E*_. Employing an MR-protocol with multiple-echo-times can be used to estimate the axonal water fraction at *TE* = 0 through direct fitting of the *T*_*E*_-dependent two-compartment model (e.g. (Veraart et al., 2018)) or through fitting to an analytic expression (Gong et al., 2020). Although physically motivated, it takes substantially longer to acquire the DWI-data than in case of a single *T*_*E*_ acquisition. Here we suggest an alternative whereby an additional scaling calibration could be used to convert model estimates into true volume fractions using standard DWI at a single *T*_*E*_. This hypothesis has to be tested by comparison to a gold standard. Most commonly, electron microscopy has served as the gold standard. But due to low contrast in EM, unmyelinated axons are difficult to discriminate from other entities such as, e.g. glial cells, and thus they are typically not quantified (Lee et al., 2019; West et al., 2018b; Zaimi et al., 2018). Therefore, the application of EM as gold standard for testing DWI-based metrics of the axonal volume also revives the question whether DWI-based *f*_*A*_-estimates are covering the whole population of axons, i.e. myelined and unmyelinated axons, or whether they are affected by the fraction of myelinated axons, only (Beaulieu & Allen, 1994).

In this study, we investigate whether additional calibration through an offset accounting for the fraction of unmyelinated axons, or a scaling factor accounting for compartmental *T*_2_ differences, or a combination of both parameters, can improve the one-to-one correspondence between DWI-based metrics of the axonal volume fraction and their corresponding electron-microscopy counterparts. To this end, we use a multi-modal, ex-vivo dataset of DWI- and EM-data of mouse WM from the corpus callosum and fornix. The dataset includes several mouse models with varying degrees of myelination ranging from hypo- to hypermyelination (Kelm et al., 2016; West et al., 2018a), thereby providing a broad dynamic range of axonal and myelin volume metrics. The EM-based *f*_*A*_ and myelin volume fraction (*f*_*M*_) are used as gold standards and utilised to validate the accuracy of DWI-based *f*_*A*_ markers derived from four different biophysical model of the axonal water fraction, WMTI, NODDI, NODDI-DTI and JESP± (Fieremans et al., 2011; Zhang et al., 2012; Edwards et al., 2017; Jespersen et al., 2018).

## 2. Background

We model WM tissue as being composed of three distinct, non-overlapping compartments quantified by the axonal (*f*_*A*_), the myelin (*f*_*M*_), and the extracellular volume fraction (*f*_*E*_) with

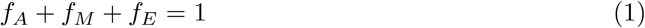

in every WM voxel (see also Fig. 1 for a schematic description of the modelled volume fractions and the metrics glossary in table 1). Since the myelin compartment is in practice not affecting the DWI-signal due to its short relaxation time, the DWI-signal is determined by the axonal water fraction (*f*_*AW*_). In the two-compartment model of WM shown in th second row of Fig. 1 *f*_*AW*_ is given by

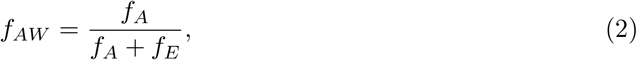

which can be converted into the axonal volume fraction *f*_*A*_ by rescaling the right side of Eq. 2 by 1 − *f*_*M*_, with *f*_*M*_ being estimated using another non-DWI technique. In order to distinguish DWI-based metrics from the “true” axonal water fraction *f*_*AW*_ given by Eq. 2, we denote them in the following by *A*_*W*_ (≈ *f*_*AW*_). For testing the performance of DWI-models, a gold standard is required for comparison. Here, we used electron microscopy metrics for both, the axonal volume fraction, denoted by *α*, and for the myelin volume fraction, denoted by *μ*, as the gold standard. To avoid ambiguities about whether the calibration parameters are correcting the DWI- or the myelin-marker, our strategy was to rescale the axonal water fraction (defined by Eqs. 1 and 2) by 1 − *μ* to get a DWI-based estimate of the axonal volume fraction *A*:

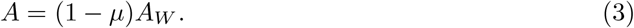

**Figure 1:**
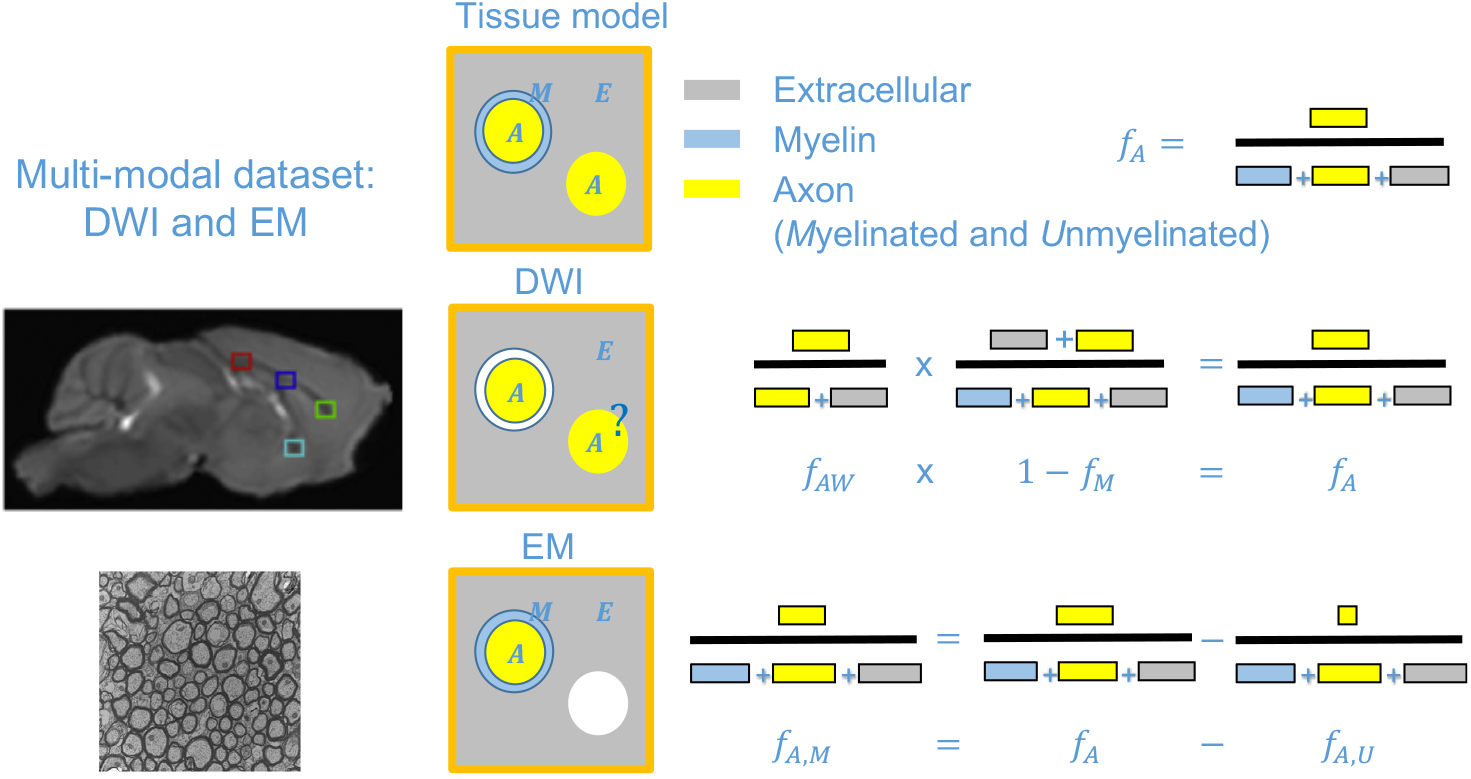
Schematic relating the tissue volume fractions of the three-compartment tissue model to their counterpart from the multi-modal dataset including diffusion-weighted-imaging- and electron microscopy-data. Note that areas presented in white are not observable with the corresponding technique, i.e. myelin in case of DWI and unmyelinated axons in case of EM. The question mark indicates that it is not known to what extent the fraction of unmyelinated axons can be estimated by the DWI models. DWI and EM images were taken from (Kelm et al., 2016) and modified. Colored boxes in the DWI image indcate regions-of-interest (ROI) in which DWI-metrics were available in this study.

**Table 1:** Summary of the employed DWI- and EM-metrics and their relation with the tissue compartment model volume fractions.

| Description | Tissue model metric | DWI metric | EM metric |
| --- | --- | --- | --- |
| Axonal volume fraction | $f_A$ | $A$ | - |
| Axonal volume fraction (unmyelinated) | $f_{A,U}$ | - | - |
| Axonal volume fraction (myelinated) | $f_{A,M}$ | - | $\alpha$ |
| Axonal water fraction | $f_{AW}$ | $A_W$ | - |
| Myelin volume fraction | $f_M$ | - | $\mu$ |
| Extracellular volume fraction | $f_E$ | - | - |

In EM only the myelinated axons are assessed, i.e.

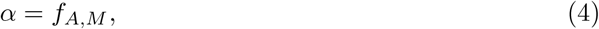

with *f*_*A,M*_ + *f*_*A,U*_ = *f*_*A*_, where the subscripts *M* and *U* denote myelinated and unmyelinated, respectively. Taking *α* (Eq. 4) as the gold standard therefore raises the question how sensitive the DWI-based signal is to the fraction of unmyelinated axons, which would occur as an additional offset calibration parameter *U* to *A*:

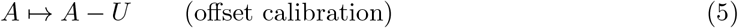

Eq. 3 is based on the assumption that apart from measurement error *A*_*W*_ is equal to *f*_*AW*_, which is a common assumption implying that compartmental *T*_2_ differences in transverse relaxation (*T*_2_) are negligible. If compartmental differences cannot be neglected, the fraction of the axonal signal would become a function of the employed echo time *T*_*E*_ with *A*_*W*_ (*T*_*E*_ = 0) = *f*_*AW*_ (see, e.g. (Gong et al., 2020)). We will show in the subsequent section that this *T*_*E*_-dependence can be separated into a scaling calibration factor:

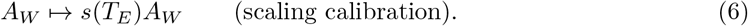

In the following we tested the hypothesis that calibration parameters *U* and *s* (Eqs. 5 and 6) can improve the one-to-one correspondence between *α* and *A* (Eqs. 3 and 4). For further details on calibration parameters and optimisation see the corresponding methods section.

### 2.1 Analytical derivation of the scaling calibration

Each of the tested DWI-models (WMTI, NODDI, NODDI-DTI, and JESP) includes a number of assumptions about the underlying WM microstructure (see, e.g. (Mohammadi & Callaghan, 2021)). All signal models tested in this study can be derived on the basis of the four-compartment ex-vivo NODDI-model composed of an axonal (with index *a*), extracellular, isotropic (*iso*) and a so-called dot (*dot*) compartment, which accounts for water trapped inside small cavities in fixed tissue with effectively no diffusivity (Alexander et al., 2010; Panagiotaki et al., 2012), given by

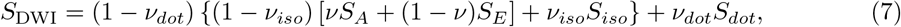

where *ν*_*dot*_, *ν*_*iso*_, and *ν* are the signal fractions of the dot-, isotropic-, and axonal compartments, respectively. The usual in-vivo NODDI-model is obtained by setting *ν*_*dot*_ = 0 and the signal models for WMTI, JESP+/- and NODDI-DTI assume *ν*_*dot*_ = *ν*_*iso*_ = 0. The compartmental signals (*S*_*dot*_, *S*_*iso*_, and *S*_*a*_) are functions of the diffusion b-vector **b** and a set of biophysical parameters {*p*_*i*_}, which depend on the DWI-model (see table 2), with the assumption that at **b** = 0 *S*_*A*_ = *S*_*E*_ = *S*_*iso*_ = *S*_*dot*_ = 1. In analogy to the volume fractions from the introduction (see also Fig. 1), the signal fractions in Eq. 7 are: *ν* = *f*_*A*_*/*(*f*_*A*_ + *f*_*E*_), *ν*_*iso*_ = *f*_*iso*_*/*(*f*_*A*_ + *f*_*E*_ + *f*_*iso*_), and *ν*_*dot*_ = *f*_*dot*_*/*(*f*_*A*_ + *f*_*E*_ + *f*_*iso*_ + *f*_*dot*_), so that *ν* + *ν*_*iso*_ + *ν*_*dot*_ = 1. In this signal model the axonal water fraction is given directly by the corresponding coefficient of *S*_*A*_: (ex-vivo NODDI) *A*_*W*_ = (1 − *ν*_*dot*_)(1 − *ν*_*iso*_)*ν*, (NODDI, i.e. *ν*_*dot*_ = 0) *A*_*W*_ = (1 − *ν*_*iso*_)*ν*, and (WMTI, JESP+/- and NODDI-DTI, i.e. *ν*_*dot*_ = *ν*_*iso*_ = 0) *A*_*W*_ = *ν*. This changes if the compartmental signals are functions of the corresponding compartmental transverse relaxation times. A simple signal model accounting for compartmental relaxation times is given by

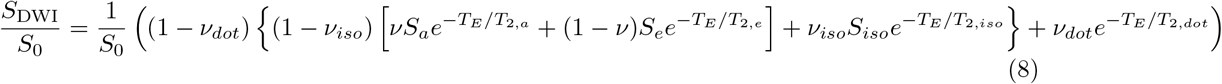

where *T*_*E*_ is the echo-time, and *T*_2,*a*_, *T*_2,*e*_ and *T*_2,*iso*_ are the transverse relaxation times in the axonal-, extra-cellular, and isotropic compartments. For this signal model *S*_0_ 1 and hence the coefficient of *S*_*a*_ is now 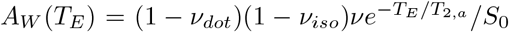 and hence *T*_*E*_-dependent. The application of a model that does not account for compartmental relaxation to diffusion MR-data will therefore require a calibration scaling factor *s*(*T*_*E*_) in order to retrieve the desired axonal water fraction from the coefficient of the axonal signal. As can be directly seen from the ex-vivo NODDI signal model Eq. 8

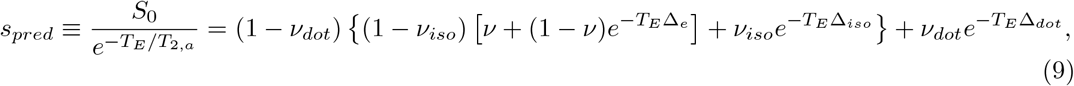

where Δ_*e*_ = 1*/*(*T*_2,*a*_ − 1*/T*_2,*e*_), Δ_*iso*_ = 1*/*(*T*_2,*a*_ − 1*/T*_2,*iso*_), and Δ_*dot*_ = 1*/*(*T*_2,*a*_ − 1*/T*_2,*dot*_). Then we have *s*_*pred*_*A*_*W*_ = (1 − *ν*_*dot*_)(1 − *ν*_*iso*_)*ν* ≡ *f*_*A*_ again. For the usual NODDI signal model *ν*_*dot*_ = 0 and for the remaining two-compartment models of this study *ν*_*dot*_ = *ν*_*iso*_ = 0. The scaling can be predicted for the DWI-models using Eq. 9 once the compartmental signal fraction, the compartmental *T*_2_ and the echo-times are known from literature or estimated from multi-echo measurements.

**Table 2:** Summary by DWI-model of the input data, the free biophysical parameters *p*_*j*_ and the assumptions on them for the four validated DWI-models. The basic signal expression is given in Eq. 7. Symbols are as follows: (*D*∥) parallel diffusivity, (*D*_⊥_) perpendicular diffusivity, (*W*∥) parallel kurtosis, (*W*_⊥_) perpendicular kurtosis, (⟨*W*⟩) mean kurtosis, (*FA*) fractional anisotropy, (*MD*) mean diffusivity, (*d*_*e*,_ ∥, *d*_*e*,⊥_) diffusivities in the extracellular compartment, (*d*_*a*,_ ∥) and in the axonal compartment, (*κ*) fibre dispersion, (*ν*) axonal signal fraction, (*ν*_*iso*_) and (*ν*_*iso*_) signal fractions of the isotropic and dot compartment, repectively.

| DWI-model | Input parameters | Biophysical parameters $p_j$ | Assumptions |
| --- | --- | --- | --- |
| WMTI | All standard DKI | $d_{e,\parallel}, d_{e,\perp}, d_{a,\parallel}, \nu$ | $\nu_{dot} = \nu_{iso} = 0, \kappa \rightarrow \infty$ |
| JESP+/- | $D_{\parallel}, D_{\perp}, \langle W \rangle, W_{\parallel}, W_{\perp}$ | $d_{e,\parallel}, d_{e,\perp}, d_{a,\parallel}, \kappa, \nu$ | $\nu_{dot} = \nu_{iso} = 0$ |
| NODDI | DWI-data | in-vivo: $\kappa, \nu, \nu_{iso}$<br>ex-vivo: $\kappa, \nu, \nu_{iso}, \nu_{dot}$ | $d_{e,\parallel} = d_{a,\parallel},$<br>$d_{e,\perp} = (1 - \nu)d_{e,\parallel},$<br>$d_{iso} = 2.0\mu^2/ms,$<br>$d_{e,\parallel} = 0.35\mu^2/ms$ |
| NODDI-DTI | $FA, MD$ | $\kappa, \nu$ | $\nu_{dot} = \nu_{iso} = 0,$<br>$d_{e,\parallel} = d_{a,\parallel},$<br>$d_{e,\perp} = (1 - \nu)d_{e,\parallel},$<br>$d_{e,\parallel} = 0.35\mu^2/ms$ |

#### 2.1.1. Estimation of compartmental T_2_ from the literature

For estimating *s*_pred_, we rescaled the compartmental *T*_2_-values *T*_2,*a*_(3*T*) ≈ 83 ms and *T*_2,*e*_(3*T*) ≈ 59 ms from (Tax et al., 2021) from 3T to 15.2T. The scaling was estimated from average values for the transverse relaxation time in ex-vivo human white matter on the basis of monoexponential models: *T*_2_(3*T*) = 83.8 ms (Birkl et al., 2014), and *T*_2_(15.2*T*) ≈ 33 ms (from the supplementary of (West et al., 2018b)). The decrease of the relaxation time from 3*T* to 15.2*T* can then be estimated by the ratio

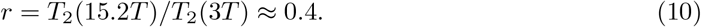

With Eq. 10 the relaxation rates of the individual compartments can be estimated as

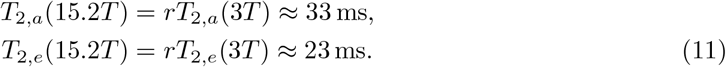

From Eqs. 11 the compartmental differences are then Δ_*e*_ ≈ 12.6 s^−1^ and Δ_*iso*_ ≈ −32.0 s^−1^, where *T*_2,*iso*_ = 1 ms was assumed (Gong et al., 2020). For *T*_2,*dot*_ no values could be found to the best of our knowledge. We assumed *T*_2,*dot*_ = *T*_2,*e*_, i.e. Δ_*dot*_ = Δ_*e*_ in Eq. 9.

#### 2.1.2. Estimation of the scaling based on theory and literature

Taking Δ_*e*_, Δ_*iso*_, and Δ_*dot*_ from the previous subsection, taking *T*_*E*_ = 19ms from (Kelm et al., 2016), and taking *ν* = 0.5 and *ν*_*iso*_ = 0.05 from Gong et al. (2020), Eq. 9 yields for NODDI *s*_*pred*_ ≈ 0.91. The predicted scaling for the remaining DWI-models (WMTI, JESP+/-, and NODDI-DTI) yields *s*_*pred*_ ≈ 0.883, following again from Eq. 9 with the assumption *ν*_*dot*_ = *ν*_*iso*_ = 0, *ν* = 0.45 from Veraart et al. (2018), and any missing constants as for NODDI.

## 3. Methods and Materials

### 3.1. Dataset

The dataset used in this study is described in detail in (Kelm et al., 2016; West et al., 2018a). The data included DWI and EM-histology data in an ex-vivo cohort of *N* = 15 mice. Six of which were healthy controls (i.e. *N*_*Controls*_ = 6) and nine were genetically modified mouse models: three Pten CKO (hypermyelinated), three Rictor CKO (hypomyelinated) and three Tsc2 CKO (severely hypomyelinated), i.e. (*N*_*Pten*_ = *N*_*Rictor*_ = *N*_*T sc*2_ = 3). In each mouse model, DWI-based axonal water fraction estimates acquired at 15.2T were available from four DWI-models (*A*_*W*_ with ∈ {DWI WMTI, NODDI, NODDI-DTI, JESP+/-}) and EM metrics *α* and *μ* in four ROIs, three in the corpus callosum (genu, mid-body, splenium) and one ROI in the fornix as indicated by the colored boxes in Fig. 1. In total, this resulted in *N* = 60 numerical values (15 mice x 4 ROIs) for each EM and DWI metric. Resolution in DWI was 150 × 150 *μm*^2^, i.e. 22500 *μm*^2^ cross-sectional voxel area. EM-based tissue metrics were assessed on images of size 2304 × 1888 px (controls) and 2048 × 1632 px (Pten, Rictor and Tsc2) at a resolution of 0.004 × 0.009 *μm*^2^, i.e. with a total area of ≈ 156 *μm*^2^ or ≈ 126 *μm*^2^, respectively. Eight data points violating the DWI-model conditions of NODDI-DTI and JESP+/- were excluded from all further analyses, i.e. all analyses were based on the remaining *N* = 52 data points. All DWI-models assume that WM fibres are impermeable sticks with no diffusion perpendicular to their orientation (Novikov et al., 2018b, 2019). EM-section size ranged between ≈ 10 × 10 to 40 × 40*μ*m^2^ with 0.022*μ*m thickness, while DWI resolution was 150 × 150 × 150*μ*m^3^.

For a summary of the biophysical output parameters of the DWI-models see table 2. All DWI-models except NODDI took as input all or a subset of the 21 standard diffusion-kurtosis-imaging (DKI) parameters, i.e., the five independent elements of the diffusion tensor and 16 independent elements of the kurtosis tensor. The DKI-parameters were estimated using the non-linear least squares DKI-framework implemented in the ACID toolbox (https://www.diffusiontools.com).

WMTI: *A*_*W*_ was estimated using the WMTI model (Fieremans et al., 2011) implemented in (https://github.com/m-ama/DKI-Designer) using all 21 DKI-parameters as input. Free biophysical parameters of WMTI are the compartmental diffusivities and the axonal water fraction. WM fibres are assumed to be in parallel.

JESP+/JESP-: *A*_*W*_ was estimated using an in-house fitting algorithm implementation of the biophysical model introduced by (Jespersen et al., 2018) including two branches (JESP+ and JESP-). Input parameters were the following five DKI-parameters: parallel diffusivity *D*, perpendicular diffusivity *D*_⊥_, parallel kurtosis *W*, perpendicular kurtosis *W*_⊥_, and mean kurtosis ⟨*W⟩*. Free parameters of the JESP DWI-model are like for WMTI but in addition also axially symmetric fibre dispersion (*κ*) is accounted for via the Watson distribution.

NODDI: *A*_*W*_ was estimated from the ex-vivo NODDI model (Zhang et al., 2012) implemented in the NODDI MATLAB toolbox (https://www.nitrc.org/projects/noddi_toolbox). In contrast to the other DWI-models, NODDI took as input directly the DWI-signal. Moreover, for the ex-vivo case NODDI features foru tissue compartments: axonal, extra-cellular, isotropic, and dot compartment. NODDI accounts for axially symmetric fibre dispersion but fixes the diffusivities of the compartments as follows: parallel diffusivities in the axonal and the extracellular spaces are the same, and the perpendicular diffusivity of the extracellular compartment is related to the parallel diffusivity via a tortuosity assumption (Szafer et al., 1995). Furthermore, the fixed parallel diffusivity (0.35 *μm*^2^ms^−1^) was chosen so that the correlation between MRI and histology was maximised (West et al., 2018a).

NODDI-DTI: *A*_*W*_ was determined from the aforementioned DKI fit using the fractional anisotropy (*FA*) and mean diffusivity (*MD*) from the standard DKI model as input. The *FA* and *MD* maps used as input for NODDI-DTI were calculated from the DKI fit as recommended in (Edwards et al., 2017) to avoid the kurtosis bias in *MD*. Model assumptions are the same as for NODDI but with *ν*_*iso*_ = *ν*_*dot*_ = 0.

#### 3.1.1. Statistical analysis of differences between the EM-based metrics of mouse models

We assessed differences in *α* across the mouse-models in terms of an ANOVA with the Null hypothesis that the mean value of *α* was the same across all models.

### 3.2. Calibration parameter combinations

While the volume fraction of unmyelinated axons *f*_*A,U*_ was not assessed with EM histology, it probably contributes to the DWI-based estimate of the axonal water fraction (Beaulieu & Allen, 1994; Sepehrband et al., 2016). To test this hypothesis it was assumed that the DWI-based estimate for the axonal water fraction was accurate, i.e. *A*_*W*_ = *f*_*A*_ for all DWI-models. This defined our baseline. We pooled the 15 individual mice and four ROIs (excluding the eight invalid data points) into two groups according to the results from an ANOVA: (group 1: comprising healthy controls and only moderately hyper- or hypomyelinated mice) controls, Pten, Rictor and (group 2: heavily hypomyelinated mice) Tsc2, respectively, since we observed only a significant difference in the EM-based axonal volume fraction *α* between these two groups but not between any of the mouse models in the first group. Then we allowed for the estimation of individual average values of the fraction of unmyelinated axons in each of the two groups (1: Controls, Pten, Rictor, and 2: Tsc2). For the purpose of optimisation they were written as column vectors **U**_*j*_ = *U*_*j*_**e**_*j*_ (with *U*_*j*_ being the mean fraction of unmyelinated axons of group *j* and **e**_*j*_ being a *N*_*j*_ × 1 vector of ones) with *j* ∈ {1, 2}, *N*_1_ = 41, and *N*_2_ = 11 (for all DWI models). In this notation Eq. 3 including an offset calibration (Eq.5) and scalar scaling (Eq.6) calibration reads

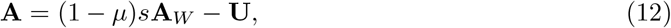

where **U** = [**U**_1_, **U**_2_]^*T*^, and *μ* and **A**_*W*_ indicate column vectors whose components are sorted in agreement with **U** and *s* is the scaling calibration parameter. To assess the sensitivity of DWI-based axonal metrics to the fraction of unmyelinated axons either a single group was allowed to vary while the other was fixed to zero, i.e. *U*_1_ ≠ 0 and *U*_2_ ≠ 0 or *U*_1_ ≠ 0 and *U*_2_ ≠ 0, and also both groups were allowed to vary individually at the same time, i.e. *U*_1_ ≠ *U*_2_ ≠ 0. For each DWI-model, we analysed six combinations of the calibration parameters, denoted in the following as {*U*_1_}, {*U*_2_}, {*s*}, {*U*_1_, *U*_2_}, {*U*_1_, *s*}, {*U*_2_, *s*}, {*U*_1_, *U*_2_, *s*}, and the baseline {}.

### 3.3. Calibration parameter estimation

The volume fractions of unmyelinated axons *U*_*j*_ and the scaling parameter *s* were estimated by minimising the residual-sum-of-squares (*RSS*) between the DWI-based estimate for the axonal volume (Eq. 12) and the EM-based gold standard (defined in Eq. 4):

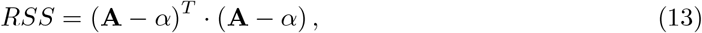

where again the bold-faced quantities represent vectors including all available numerical values assembled into column vectors. In order to ensure physically reasonable estimates of the axon volume fraction, the optimization function 7 had to be complemented by two boundary conditions. The first constraint concerns the upper limit of the sum of volume fractions, i.e., *A*+*μ* ≤ 1. The second set of constraints is concerned with the scaling parameter *s* (Eq. 6), which is required to yield values larger or equal to zero, i.e., *sA*_*W*,DWI_ ≥ 0. All parameter estimations were performed using Matlab 2020a (Mathworks, CA, USA).

### 3.4. Calibration parameter selection

To quantify the intra-model performance improvement of each DWI-model due to the calibration parameters *U*_*j*_ and *s*, we used the Bayesian information criterion (*BIC*) (Burnham et al., 2002):

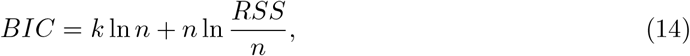

where *k* is the number of model parameters, which varied between zero (baseline {}) and three depending on the combination of *U*_*j*_, *n* is the number of evaluated data points, and *RSS* is defined in Eq. 13. The *BIC* measures a model’s capability of explaining given data while penalising overfitting. A lower *BIC* indicates less information loss, meaning that the model with the lowest *BIC* best explains the data. Since we employed the uncalibrated case {} as baseline, we only report differences Δ*BIC* with respect to this case, i.e.

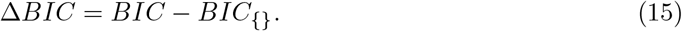

Δ*BIC* was always calculated using all available data. For assessing the variation in *U*_*j*_ and *s*, we performed a leave-one-out analysis by successively discarding the data of one mouse individually until each mouse was left out once.

### 3.5. Correspondence to EM data

To asses the models’ capability to explain the EM-based gold standard, we performed a Bland-Altman (BA) analysis (Martin Bland & Altman, 1986) of the differences

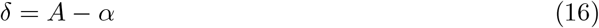

versus the mean

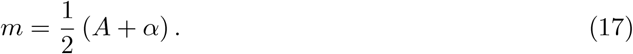

As usual, the error given by

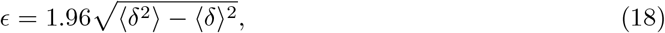

is reported. Furthermore, we also report the mean difference ⟨*δ*⟩, i.e. the bias, and bias 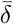 and error 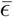 relative to the dynamic range in the EM-based axonal metric, i.e.

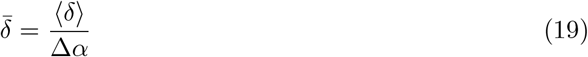

and

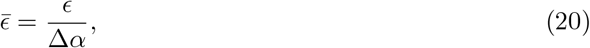

where Δ*α* = *α*_*max*_ − *α*_*min*_ and angles indicate average.

## 4. Results

We tested the hypothesis that linear calibration including a scaling parameter *s* accounting for compartmental relaxation, and an appropriate number of offsets *U*_*j*_ accounting for the volume fraction of unmyelinated axons in the different groups of mice (see Eq. 12), can improve the accuracy of DWI-based axonal volume estimates of four DWI-models (WMTI, JESP+/-, NODDI, and NODDI-DTI). Furthermore on the basis of a simple DWI-signal expression including compartmental *T*_2_ (Eq. 8), we predicted the scaling calibration (Eq. 9) and analysed how much improvement can be achieved if the scaling is fixed to the predicted values and only the offset is estimated on the basis of data. Our analyses can be divided into four steps.

First, we statistically assessed differences of the mouse models with respect to the EM-based axonal volume *α*. The maximum number of mouse models that could be discriminated statistically corresponded to the maximum number of offset parameters that were used in the subsequent analyses.

In a second step, we determined via data fitting, which combination of calibration parameters has the highest evidence of improving the one-to-one correspondence between DWI-based axonal volume *A* and its EM-based counterpart *α*. To this end we employed the Bayesian Information Criterion (*BIC* in Eqs. 14 and 15).

Third, we fixed the scaling parameter of the best calibration parameter combination determined in the previous analysis to the value predicted by our theory and only determined the remaining offset as in the second step via a fit to the data.

Finally, we compared the accuracy of *A* for all DWI-models in terms of relative bias and error (Eqs. 16 - 19) between the axonal volume fraction from DWI, *A*, and EM, *α*, for three combinations of calibration parameters: (i) the baseline with no additional calibration, (ii) the best combination determined in the second step, and (iii) the best combination but with the scaling fixed to the predicted value.

### 4.1. Statistical analysis of differences between the EM-based metrics of mouse models

To prevent subsequent parameter optimisation from modeling noisy data, we performed an ANOVA on the EM-based gold standard axonal volume fraction *α*. For this analysis, we assumed that the total axonal volume fraction, i.e. the sum of unmyelinated and myelinated axons, is similar across all mouse models and thus the measured differences in *α* essentially reflect the differences in *f*_*A,U*_. The result of the analysis shown in Fig. 2 revealed a significant (*p <* 0.05) difference between the Tsc2 mice and any of the other models, while no significant differences were observed among Pten, Rictor and Controls. As a consequence, the four mouse models were pooled into two groups, (1) controls, Pten, Rictor, and (2) Tsc2 for further analysis. For each group, one offset parameter correcting for the fraction of unmyelinated axons was introduced: *U*_1_ for group 1 and *U*_2_ for group 2.

**Figure 2:**
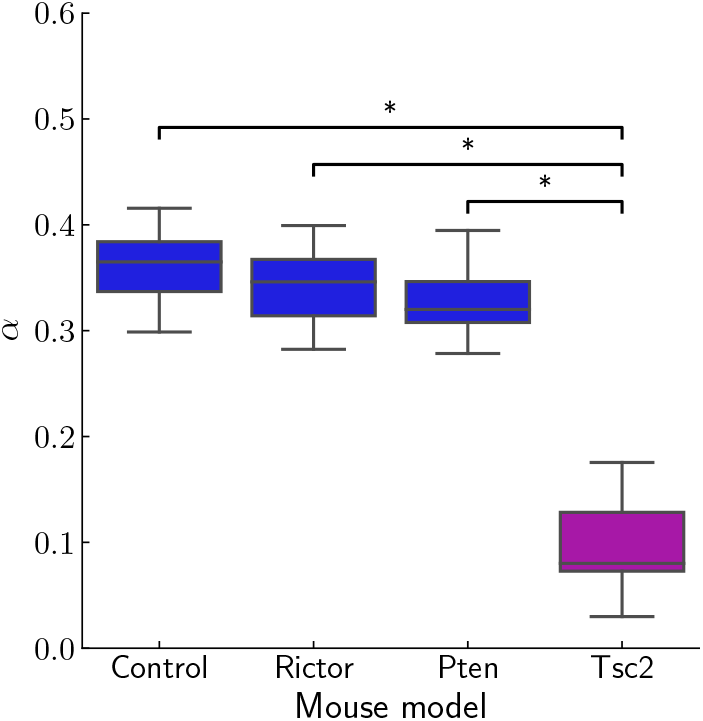
EM-based axonal volume fraction *α* for the four mouse models controls, Pten, Rictor and Tsc2. An ANOVA revealed significant differences (*p <* 0.05) only between Tsc2 and Rictor, controls or Pten, respectively. No further significant differences were observed. This motivated the pooling of the data into two groups: (1) controls, Rictor, and Pten, and (2) Tsc2.

### 4.2. Calibration parameter selection

For selecting the best combination of the offset calibration parameters, *U*_1_ and *U*_2_ for the two groups of mice, and the scaling factor *s*, we tested all possible combinations of calibration parameters. As optimisation function we used the residual-sum-of-squares between *A* and *α* (Eq. 13). To evaluate each model we determined the *BIC* (Eq. 14) and report the differences Δ*BIC* (Eq. 15) with respect to the uncalibrated case (baseline). The Δ*BIC* for the tested parameter combinations are shown in Fig. 3 for each DWI model separately. Greatest evidence for improvement was achieved in most of the DWI-models for the {*U*_2_, *s*} combination of calibration parameters, i.e. when the fraction of unmyelinated axons for the severely hypomyelinated group 2 (*U*_2_) was included together with the scaling (*s*), except for JESP- and NODDI, for which {*U*_2_} and {*U*_2_, *s*} performed equally well.

**Figure 3:**
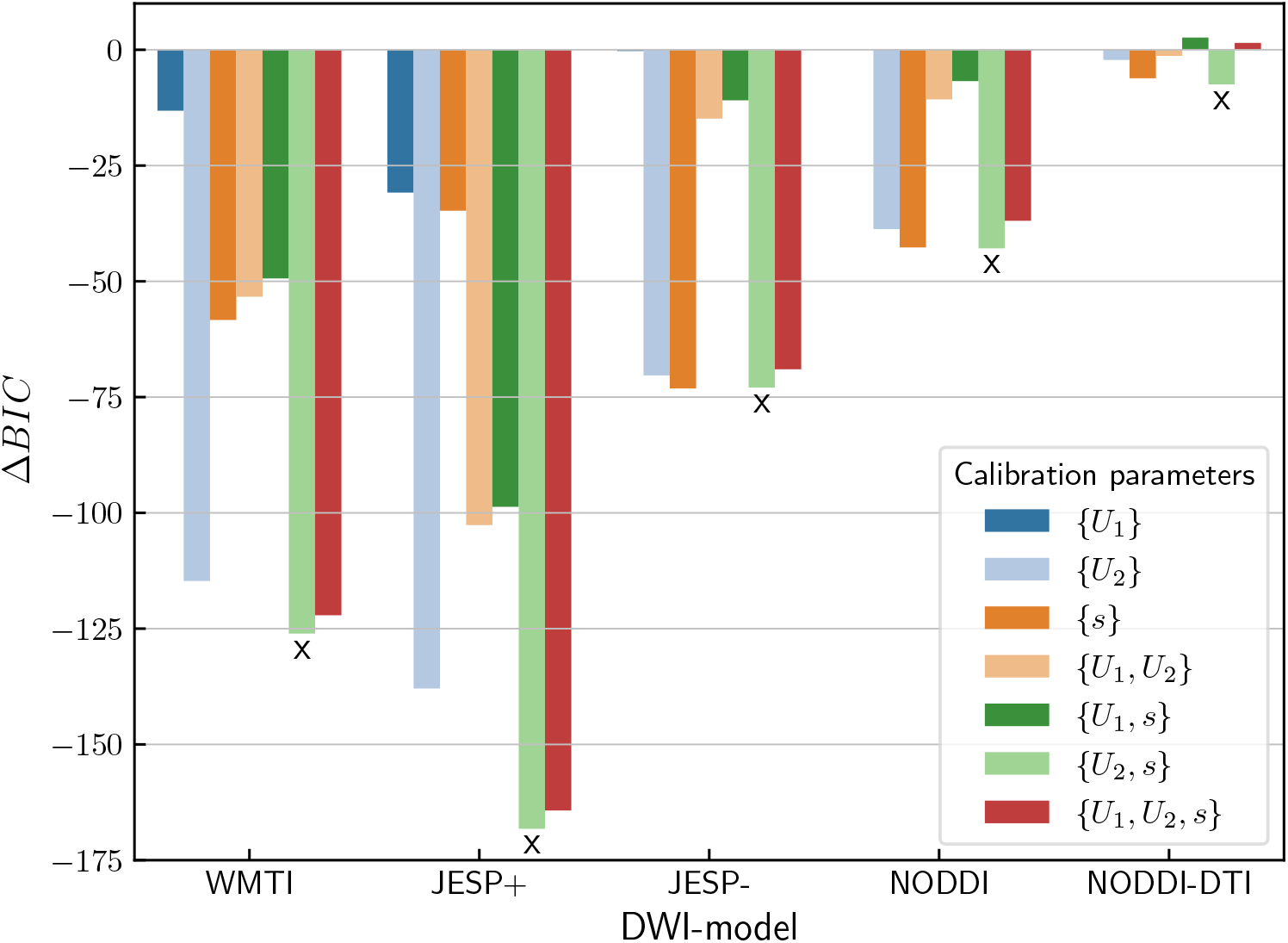
Contribution of calibration parameters to DWI-model improvement. Shown are the differences Δ*BIC* (Eq. 15) with respect to the parameter combination with smallest *BIC* in each DWI-model. A lower value indicates better model performance. Values for the uncalibrated case *{}* served as baseline, i.e. for this case Δ*BIC* = 0. For the individual *BIC* see also table 4. An x indicates the calibration parameter combination with the largest evidence of improvement with respect to the baseline without further calibration for each DWI-model, respectively.

#### 4.2.1. Best performing calibration parameter combinations

Table 4 summarises the *BIC*, volume fractions of unmyelinated axons *U*_2_, scaling *s* as determined based on data. Values are given the for the baseline ({}) without calibration and for the best combination of calibration parameters as indicated in Fig. 3. For a summary of all validated calibration parameter combinations see table A.6 in the appendix. The estimated average volume fractions of unmyelinated axons *U*_2_ for the Tsc2 mouse-model varied between 0.19 - 0.30 for WMTI, JESP, and NODDI, while it was much lower with 0.08 for NODDI-DTI. The scaling varied between 0.6 and 0.97 (note that a scaling *s* = 1 is equivalent to no scaling calibration).

**Table 3:** Summary of the tested calibration parameters. Groups of mice are defined in Fig. 2.

| Description | Calibration parameter |
| --- | --- |
| Estimate of $f_{A,U}$ of the first group of mice | $U_1$ |
| Estimate of $f_{A,U}$ of the second group of mice | $U_2$ |
| Scaling accounting for differences in compartmental $T_2$ | $s$ |

**Table 4:** Summary of *BIC*, volume fraction of unmyelinated axons *U*_2_, scaling *s* and relative difference between fit *s* and theory *s*_pred_, i.e. Δ*s* = (*s − s*_pred_)*/s*_pred_ for the baseline and the best parameter combinations from the first analysis (see also Fig. 3). Note that for determining *BIC* always all available data was used and hence no standard deviation (SD) was given as is common praxis (Burnham et al., 2002). In contrast, *U*_2_ and *s* were estimated in a leave-one-out fashion, in which each mouse individual was excluded from the computation once in order to get an estimate of the standard deviation.

| DWI-model | Calibr. parameters | $BIC$ | $U_2$ (SD) | $s$ (SD) | $s_{\text{pred}}$ | Rel. $\Delta s$ [%] |
| --- | --- | --- | --- | --- | --- | --- |
| WMTI | $\{\}$ | -174 | 0 (-) | 1 (-) | - | - |
| | $\{U_2, s\}$ | -300 | 0.219 (0.004) | 0.783 (0.004) | 0.883 | -11 |
| JESP+ | $\{\}$ | -120 | 0 (-) | 1 (-) | - | - |
| | $\{U_2, s\}$ | -288 | 0.226 (0.004) | 0.603 (0.003) | 0.883 | -31 |
| JESP- | $\{\}$ | -207 | 0 (-) | 1 (-) | - | - |
| | $\{U_2, s\}$ | -280 | 0.221 (0.004) | 0.949 (0.007) | 0.883 | 7 |
| NODDI | $\{\}$ | -204 | 0 (-) | 1 (-) | - | - |
| | $\{U_2, s\}$ | -247 | 0.194 (0.005) | 0.945 (0.005) | 0.91 | 3 |
| NODDI-DTI | $\{\}$ | -214 | 0 (-) | 1 (-) | - | - |
| | $\{U_2, s\}$ | -222 | 0.08 (0.004) | 0.972 (0.016) | 0.883 | 10 |

### 4.3. Prediction of the scaling calibration factor from theory

Based on a simple model of the DWI-signal including different *T*_2_-times in the compartments (Eq. 8), we derived analytical expressions of the scaling calibration for WMTI, JESP+/-, and NODDI-DTI and for NODDI (Eq. 9). Using *T*_2_ times (section 2.1.1) metrics (section 2.1.2) from the literature and we found *s*_*pred*_ ≈ 0.91 for NODDI and *s*_*pred*_ ≈ 0.883 for the other DWI-models. Table 4 shows that the smallest relative difference between fitted and predicted scaling Δ*s* was found for NODDI (3%) and the largest relative difference was found for JESP+ (−31%). Details about how the scaling was predicted are given in the corresponding methods sections.

### 4.4. Correspondence to EM data before and after calibration

We assessed the one-to-one correspondence of the DWI-based axonal volume fraction *A* to the EM-based axonal volume fraction *α* for the baseline and for the best performing calibration parameter combination {*U*_2_, *s*} from the previous analyses. Furthermore, we quantified change of the accuracy, gained by introducing the two calibration parameters, when the scaling is replaced with the value predicted by theory and only *U*_2_ was determined based on data, i.e. for the combination {*U*_2_, *s*_*pred*_}. To this end we report a Bland-Altman-analysis of the bias and variance. Moreover, we report bias and variance relative to the dynamic range (Δ*α*) observed in the gold standard (see Eq. 18 and Eq. 19).

In Fig. 4 we compare different scatter-plots of the histological gold standard *α* versus its DWI-based counterpart *A* for the calibration parameters combinations (i)-(iii) described in the introduction of the results. The first row of the figure shows the baseline (i), i.e. without any calibration parameters. WMTI and JESP+ clearly overestimated the axonal volume fraction in both groups of mice indicated by the global offset from the line of unity. For JESP- and NODDI only the Tsc2 mice featured an obvious positive offset, while for NODDI-DTI all mouse models showed considerably better one-to-one mapping, although with large variance along *A*. The second row shows the two-parameter calibration using {*U*_2_, *s*}, which performed the best for all DWI-models (ii). A substantially better one-to-one mapping was observed for all models except for NODDI-DTI, whereby WMTI and JESP+ appear to benefit the most by the additional calibration. The third row shows the scatter-plots for the calibration parameter combination {*U*_2_, *s*_*pred*_}, i.e. when the scaling was fixed to its predicted value and only the offset was determined via fit to the data (iii). A similar degree of improvement as for {*U*_2_, *s*} (second row) was observed for all models except JESP+.

**Figure 4:**
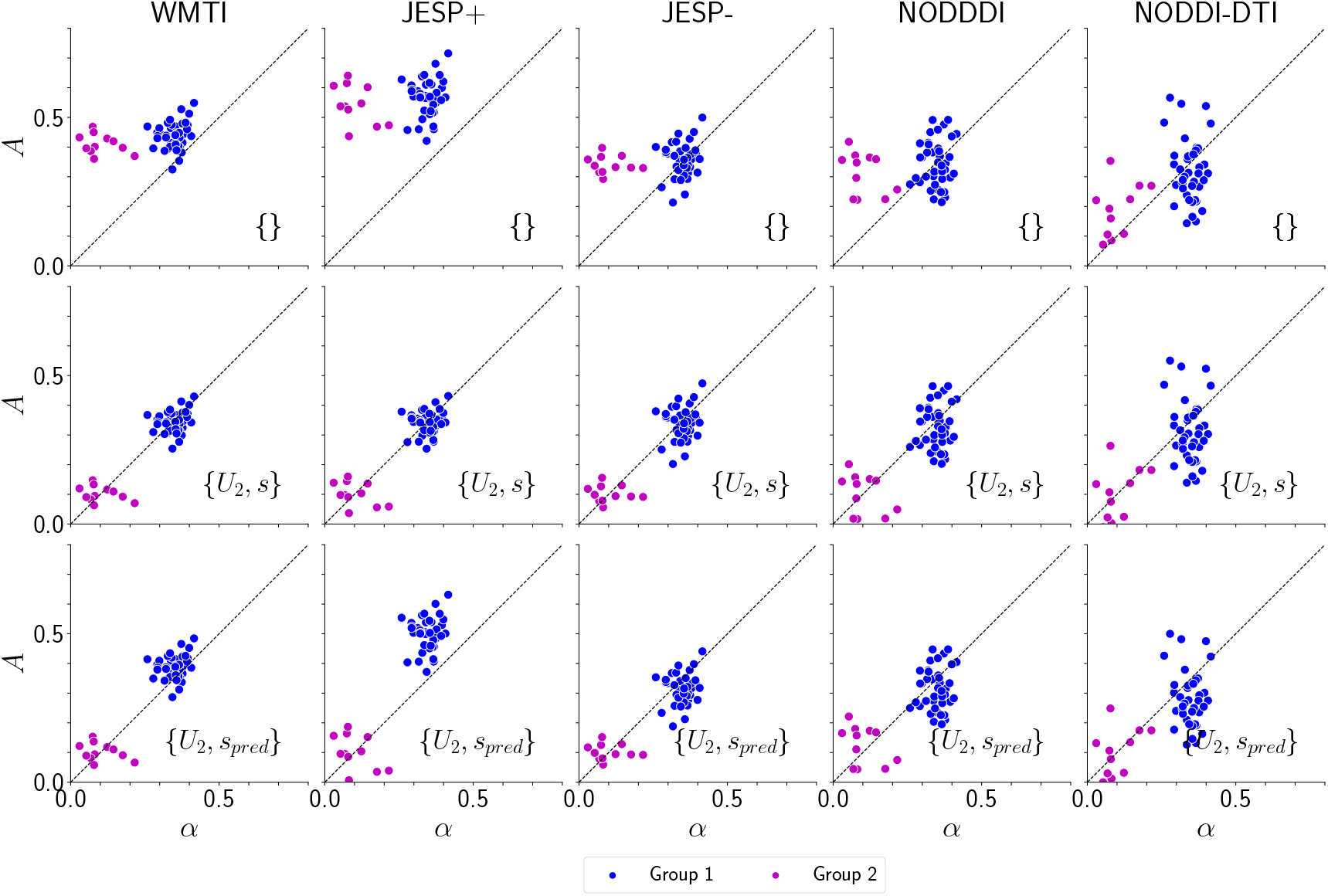
Scatter plots showing gold standard *α* (EM) vs DWI-based estimates of the axonal volume fraction (*A*). The first row shows the baseline, i.e. without additional calibration parameters and subsequent rows show the best (in terms of Δ*BIC*) calibration parameter combinations (see also Fig. 3 and table 4) with both parameters determined based on data (second row) and with the scaling fixed to the predicted values (last row). Data shown pool four regions-of-interest per mouse individual. The data were divided into two groups determined by statistical distinguishability observed in the EM gold standard *α* (see Fig. 2). The two groups are: (1) controls, Rictor, and Pten (blue), and (2) Tsc2 (magenta).)

The capability to predict the EM-based gold standard is quantified in terms of Bland-Altman (BA) plots shown in Fig. 5 and bias and error relative to the dynamic range of the EM gold standard summarised in table 5. The results for the same calibration parameter combinations (i)-(iii) as in Fig. 4 are shown. The BA-plots show a substantial reduction in bias and error for the best calibration parameter combination {*U*_2_, *s*} for all DWI-models, except for the bias of NODDI-DTI, for which the bias increased while the error decreased. Bias (error) relative to the dynamic range in the gold standard *α* was lowest for WMTI 0%(22%), followed by JESP+ 0%(24%), JESP-2%(26%), NODDI 3%(36%), and NODDI-DTI 3%(46%). When the scaling was fixed to its predicted value, the relative bias and error increased only by a few percent except for JESP+, indicating that the predicted scaling factor can be used as a proxy without substantial loss in accuracy. For JESP+ the increase was larger, but relative error and bias were still lower than in case of no additional calibration.

**Figure 5:**
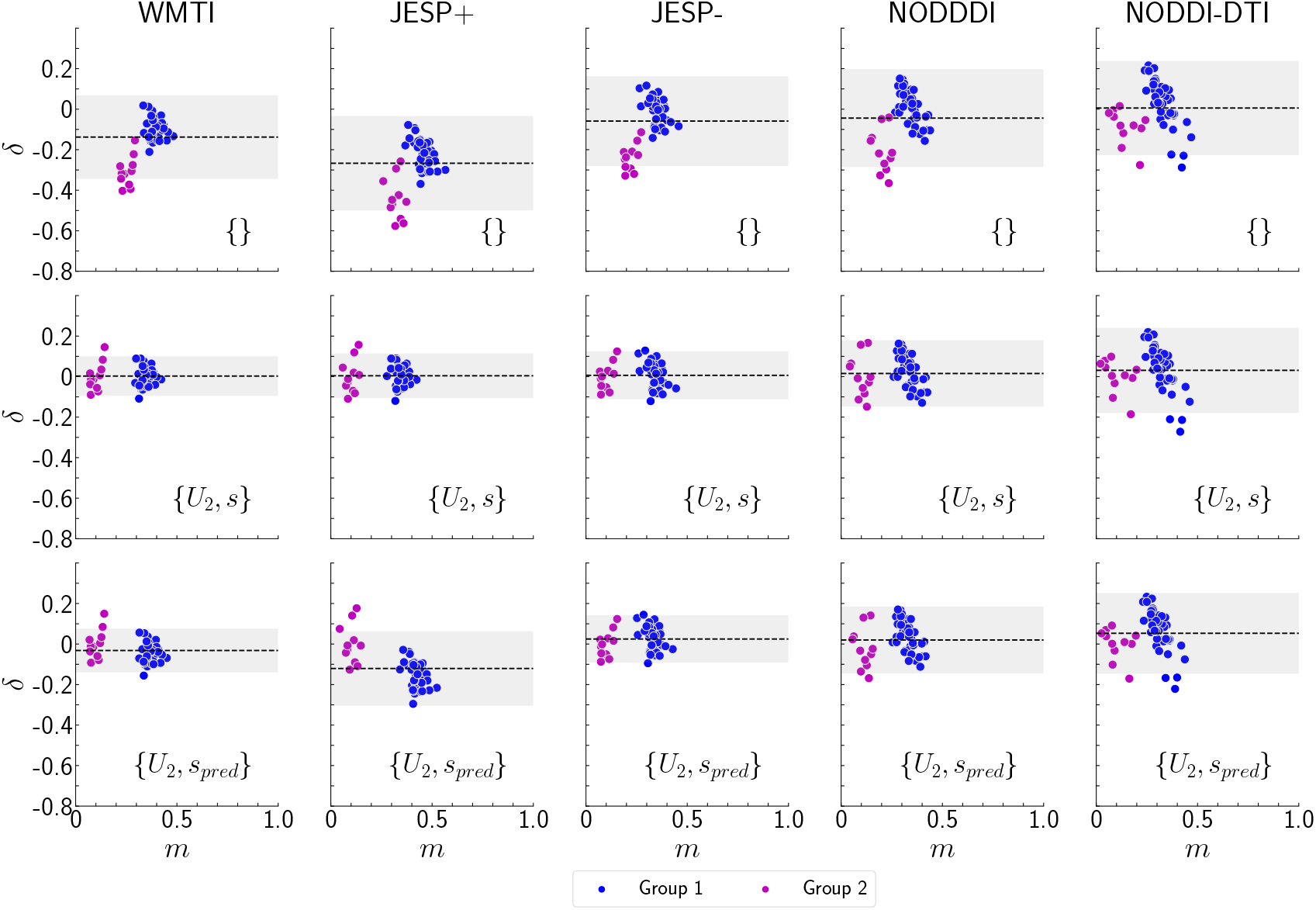
Bland-Altman plots of differences *δ* (Eq. 16) versus mean *m* (Eq. 17) between EM and DWI for the baseline (first row) and the best performing calibration parameter combination where all parameters were estimated on the basis of data (second row), and where the scaling was fixed to the theoretically predicted values (last row)(see also section 2.1.2). The dashed line corresponds to the bias ⟨*δ*⟩, while the shaded region corresponds to ⟨*δ*⟩ ± *ϵ* (see section 3.5). Individual data shown is in correspondence with Fig. 4.

**Table 5:**
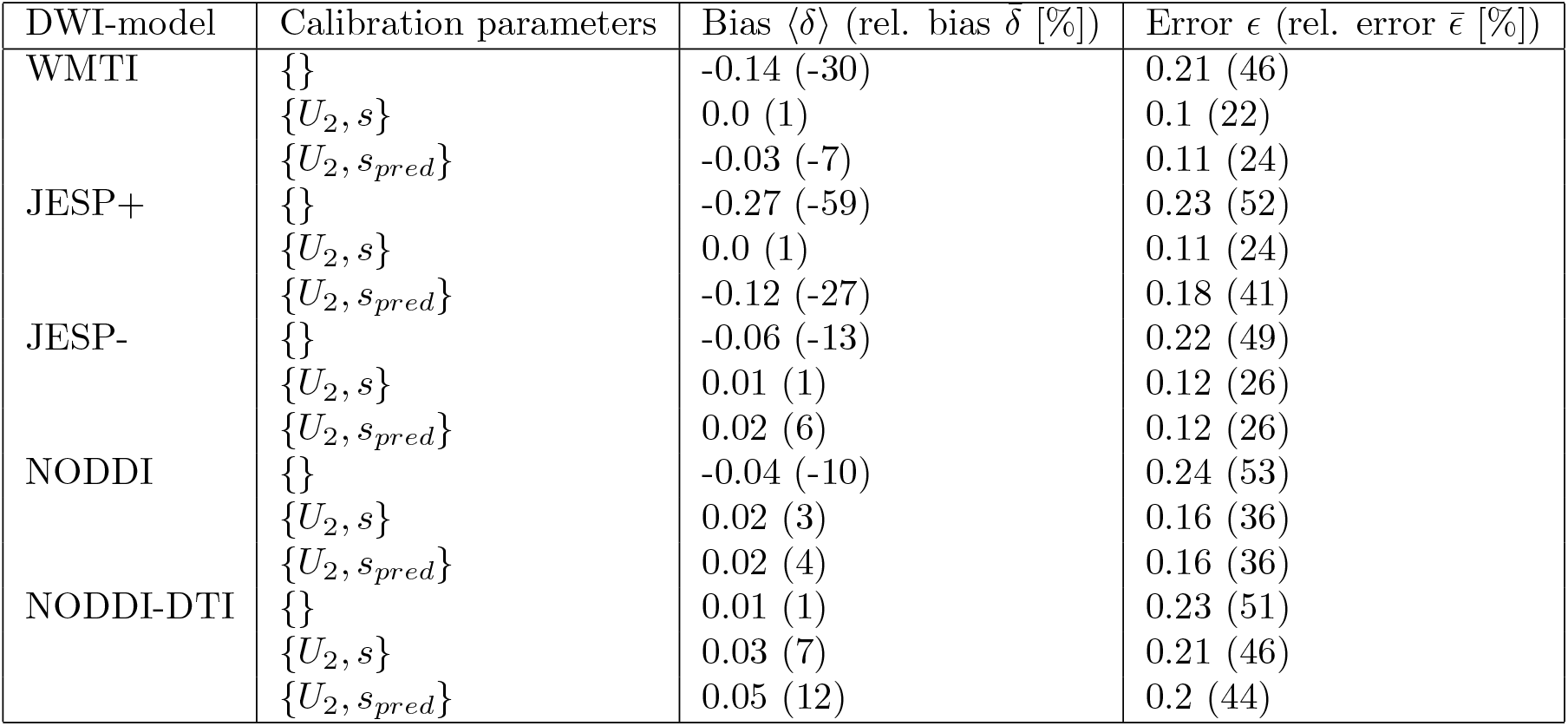
Summary of the metrics assessed to validate the capability of the re-calibrated DWI-models to predict the EM-based gold standard. Depicted are bias (relative bias) and error (relative error) for the baseline and the best performing single- and multi-parameter combinations.

## 5. Discussion

In this study we verified the hypothesis that linear calibration improves the accuracy of axonal volume fraction estimates from a range of DWI-based models with varying complexity. Using a multi-modal dataset of ex-vivo mouse models with varying degrees of myelination, we could identify two biophysically motivated calibration parameters, which substantially improve the accuracy of DWI-based axonal volume fraction estimates: one offset accounting for the fraction of unmyelinated axons of the most severely hypomyelinated mouse model, and a scaling factor accounting for compartmental differences in *T*_2_-relaxation. We propose a hybrid method to determine these two calibration parameters, in which the scaling is determined analytically and the offset is estimated from multi-modal data.

### 5.1. Calibration parameters

The biophysical interpretation of the fitted calibration parameters provides new insights about the DWI-models. First, it revealed that the sensitivity to the fraction of unmyelinated axons of investigated DWI-models is rather limited. Including the fraction of unmyelinated axons could substantially improve the model performance, but only an effect due to the most hypomyelinated mouse model could be detected (*U*_2_). For the first group of less affected mouse models the fraction of unmyelinated axons (modeled by *U*_1_) could not be detected. Furthermore, on the basis of a simple approximation, we expect that the fraction of unmyelinated axons *U*_2_ in hypomyelinated mice (ranging from 0.08 (NODDI-DTI) to 0.22 (JESP+/-)) was underestimated as well. This expectation is based on the approximation that the total axonal volume fraction (i.e. the sum of myelinated and unmyelinated axons) is the same for all mouse models although their relative proportion might change across mouse models. Given this approximation, the fraction of unmyelinated axons can be estimated as follows: observing from Fig. 2 that the fraction of myelinated axons is about 0.35 and taking the percentage (of the fraction of all axons) of unmyelinated axons reported in the literature, e.g. 33% in (Abdollahzadeh et al., 2019) or 30% in (Jelescu et al., 2016), the total axonal volume would approximately be given by ≈ 0.35*/*(1.0 − 0.33) ≈ 0.50 to 0.52. Assuming this total axonal volume fraction in Tsc2 mice (we used 0.5), we can estimate from Fig. 2 the fraction of unmyelinated axons to be 0.4, which is larger than the DWI-model-based predictions. While our results confirm the knowledge that the unmyelinated axons contribute to the DWI-signal in general (Beaulieu & Allen, 1994) (Takahashi et al., 2002) (Beaulieu, 2009), they additionally suggest a rather limited sensitivity to this fraction of axons.

Second, differences between the fitted and biophysically predicted scaling parameters divide the DWI-models into two groups: WMTI and JESP+ yield values below the theoretical value, and JESP-, NODDI, and NODDI-DTI yield values larger than predicted. In part this might be related to underlying non-uniqueness of solutions of the standard model (Jelescu et al., 2016; Novikov et al., 2018a). Hereby, WMTI and JESP+ employ the positive branch of the solutions, while JESP-represents the negative branch. In turn, NODDI and NODDI-DTI impose constraints on the relation between the axonal and the extracellular diffusivities. The intra- and extracellular diffusivities, however, determine the compartmental signals and hence constraints on them such as, e.g. fixed diffusivities or the tortuosity condition, could affect the estimated scaling. Interestingly, our findings for the scaling *s*, compared to the theoretically predicted value *s*_pred_, are in favor of the negative branch JESP-, which contradicts the common agreement in the literature that the positive branch JESP+ yields more realistic model parameters than the negative branch (Jelescu et al., 2020). This apparent contradiction could be related to the neglection of macromolecules other than in myelin in our tissue model, e.g. in the membranes of astrocytes. This means the rescaling of the axonal water fraction given by Eq. 3 should more rigorously be 1 − *μ* − *f*_*NM*_, where *f*_*NM*_ would account for non-myelin macromolecules. As a consequence, one would expect the scaling calibration parameter *s* to be smaller than the theoretical scaling parameter *s*_pred_.

### 5.2. Impact of calibration parameters to the accuracy of DWI-models

There are a number of observations related to the response of the tested DWI-models with respect to the calibration parameters and in comparison with each other that deserves further discussion. While without calibration all DWI-models are similarly inaccurate (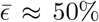, see table 5), DWI-models with fewer free model parameters improve less through calibration. While WMTI benefits the most from calibration, followed by JESP+, and JESP-, calibration had much less of an impact on NODDI, and finally NODDI-DTI hardly benefits from any of the calibration parameters. Thereby it is striking that the accuracy of the DWI-models, which fit the compartmental diffusivities, improves the most. The observation that WMTI seems to be superior to JESP was surprising because JESP also accounts for fibre dispersion whereas WMTI does not. A possible explanation could be that the larger number of free parameters in the JESP-model leads to a less well conditioned optimisation problem, which makes JESP potentially more prone to noise. The additional drop in the impact of the calibration parameters from NODDI to NODDI-DTI could be explained by the fact that NODDI is modeling two additional compartments (isotropic and dot) whereas NODDI-DTI is not. Interestingly, NODDI-DTI had the best one-to-one correspondence for the baseline, i.e. without calibration parameters, as seen in the scatter plots, which could be due to the fact that in the implementation of NODDI-DTI in the ACID toolbox (https://www.diffusiontools.com) voxels violating NODDI-DTI model assumptions are automatically identified and were removed in our study from any further analysis for all DWI-models. If only a single calibration parameter was estimated, JESP- and WMTI performed best followed by NODDI, whereas JESP+ improved less in terms of relative error and NODDI-DTI hardly showed any improvement. Here, it is also worth noting the different behaviour of the two branches JESP+ and JESP-when a second calibration parameter is included. In contrast to the positive branch, JESP-does not improve through the additional scaling. This might be due to the absence of proper minima in the optimisation function of JESP- as indicated in the original publication (Jespersen et al., 2018). Therefore, it seems recommended to treat the parameters of the JESP-branch with some caution.

### 5.3. Practical impact

Our findings have a potentially significant impact on the interpretation of existing neuroscience studies using DWI-based biophysical models to estimate the axonal volume fraction. We showed that commonly used DWI-based metrics of the axonal volume fraction, e.g. from WMTI and NODDI, possess a large error (46-53%, see table 5) if no calibration is used and thus biophysical interpretation of the results should be avoided. This is particularly important for MR-based g-ratio mapping where the axonal volume fraction from NODDI or WMTI is combined with a metric of the myelin volume fraction. Our study shows that additional calibration of the DWI-based metric can substantially improve the accuracy of axonal volume fraction estimation and thus should be employed for MR-based g-ratio mapping. Furthermore, the biophysical interpretation of the calibration parameters have practical impact. For example, we demonstrated for the standard DWI and the NODDI signal models with compartmental *T*_2_-dependence, that the scaling parameter for the calibration of the axonal volume can be predicted independently of the MRI protocol if the compartmental *T*_2_-values are known. Furthermore, using the predicted values and an offset *U*_2_ determined based on data achieved similar accuracy to estimating both calibration parameters based on EM histology data. This is of practical relevance because it allows the remaining offset calibration parameter to be estimated from a limited amount of multi-modal data and thus makes the proposed method usable in human brain data. Moreover, our finding that the contribution of the fraction of unmyelinated axons to the DWI-signal was only detectable with the investigated DWI-models for the severely hypomyelinated mouse model is of practical importance in clinical research. For example, it was suggested that it is unlikely that the changes of the axonal volume fraction observed in multiple sclerosis patients (Yu et al., 2019) are directly driven by the fraction of unmyelinated axons.

### 5.4. Limitations

Although the additional calibration parameters substantially improved the investigated DWI-models’ accuracy, the error relative to the dynamic range in the gold standard remains rather large (22% for the best model, WMTI). This remaining, rather large error may partly be attributed to a potentially large variance in the gold standard originating in the relatively small EM-section size which is probably not sufficient to representatively capture the distribution of axons in the MRI voxels (cross-sectional area of MRI-voxels was ≈ 144 (for controls) or ≈ 187 (for Pten, Rictor, and Tsc2) times larger than for EM-sections). Also the fact that our measure for the DWI-based axonal volume *A* is actually a hybrid DWI-EM-measure cannot entirely be excluded as an additional source of variance. However, according to the literature the EM-based myelin volume *μ* can be assumed to be rather accurate (West et al., 2018b)), hence unwanted variance in *A* is more likely due to the DWI-based axonal water fraction *A*_*W*_ than due to the EM-based *μ*. We favored this approach over an approach which would have involved an MRI-biomarker of the myelin volume fraction to avoid ambiguities since it would be unclear whether any observed improvement through calibration should be attributed to a correction of MRI-based myelin volume or DWI-based axonal volume. Finally, we estimated the theoretical scaling factors on the basis of in-vivo compartmental *T*_2_ values rescaled from 3T to 15.2T by a factor estimated from ex-vivo values in human brain and finally compared it to ex-vivo mouse models. Despite these limitations the predicted scaling factors *s*_*pred*_ improved the accuracy of most of the DWI-models compared to the case where the scaling is neglected (see table A.6), indicating that the chosen approach to estimate the scaling factor is valid.

### 5.5. Conclusion

In summary, we demonstrated that two biophysically motivated calibration parameters, an offset accounting for the fraction of unmyelinated axons and a scaling factor accounting for compartmental *T*_2_ differences, can improve the accuracy of four DWI-based models of the axonal volume fraction. Our findings suggest that after calibration WMTI was the most accurate of the tested DWI-models, followed by JESP+ and NODDI. NODDI-DTI hardly improved through calibration, indicating an interaction between free model parameters and efficacy of calibration. The fact that calibration improved the accuracy of the DWI-models opens the perspective to use calibrated DWI-models as an alternative to more complex biophysical models, which typically require more elaborate data acquisitions and may be prone to numerical instability. On the other hand, once determined the biophysical interpretation of the calibration parameters enables their subsequent application to more complex models as fixed model parameters.

## Acknowledgements

The research leading to these results has received funding from the European Research Council under the European Union’s Seventh Framework Programme (FP7/2007-2013) / ERC grant agreement No. 616905.

This work was supported by the German Research Foundation (DFG Priority Program 2041 “Computational Connectomics”, [MO 2397/5-1; MO 2249/3-1; GE 2967/1-1], by the Emmy Noether Stipend: MO 2397/4-1) and by the BMBF (01EW1711A and B) in the framework of ERA-NET NEURON and the Forschungszentrums Medizintechnik Hamburg (fmthh; grant 01fmthh2017).

## Appendix A. Additional data

**Table A.6:** Summary of all fitted models and calibration parameter combinations as shown in Fig. 3. The *BIC* was determined on the basis of all available data. In contrast, *U*_1_, *U*_2_ and *s* were estimated in a leave-one-out fashion, in which each mouse individual was excluded from the computation once in order to get an estimate of the standard deviation (see also table 4). The predicted scaling, *s*_*pred*_, was calculated as described in section 2.1.2.

| DWI-model | Calibr. parameters | $BIC$ | $U_1$ (SD) | $U_2$ (SD) | $s$ (SD) | $s_{pred}$ | Rel. $\Delta s$ [%] |
| --- | --- | --- | --- | --- | --- | --- | --- |
| WMTI | $\{\}$ | -174 | 0 (-) | 0 (-) | 1 (-) | - | - |
| | $\{U_1\}$ | -188 | 0.092 (0.002) | - (-) | 1 (-) | - | - |
| | $\{U_2\}$ | -233 | - (-) | 0.308 (0.004) | 1 (-) | - | - |
| | $\{U_1, U_2\}$ | -228 | 0.092 (0.002) | 0.308 (0.004) | 1 (-) | - | - |
| | $\{s\}$ | -289 | - (-) | - (-) | 0.682 (0.016) | 0.883 | -22 |
| | $\{U_1, s\}$ | -224 | 0.0 (0.0) | - (-) | 0.682 (0.016) | 0.883 | -22 |
| | $\{U_2, s\}$ | -300 | - (-) | 0.219 (0.004) | 0.783 (0.004) | 0.883 | -11 |
| | $\{U_1, U_2, s\}$ | -296 | 0.0 (0.0) | 0.219 (0.004) | 0.783 (0.004) | 0.883 | -11 |
| JESP+ | $\{\}$ | -120 | 0 (-) | 0 (-) | 1 (-) | - | - |
| | $\{U_1\}$ | -151 | 0.22 (0.003) | - (-) | 1 (-) | - | - |
| | $\{U_2\}$ | -154 | - (-) | 0.443 (0.003) | 1 (-) | - | - |
| | $\{U_1, U_2\}$ | -222 | 0.22 (0.003) | 0.443 (0.003) | 1 (-) | - | - |
| | $\{s\}$ | -258 | - (-) | - (-) | 0.521 (0.013) | 0.883 | -40 |
| | $\{U_1, s\}$ | -218 | 0.0 (0.0) | - (-) | 0.521 (0.013) | 0.883 | -40 |
| | $\{U_2, s\}$ | -288 | - (-) | 0.226 (0.004) | 0.603 (0.003) | 0.883 | -31 |
| | $\{U_1, U_2, s\}$ | -284 | 0.0 (0.0) | 0.226 (0.004) | 0.603 (0.003) | 0.883 | -31 |
| JESP- | $\{\}$ | -207 | 0 (-) | 0 (-) | 1 (-) | - | - |
| | $\{U_1\}$ | -207 | 0.01 (0.003) | - (-) | 1 (-) | - | - |
| | $\{U_2\}$ | -280 | - (-) | 0.239 (0.003) | 1 (-) | - | - |
| | $\{U_1, U_2\}$ | -222 | 0.01 (0.003) | 0.239 (0.003) | 1 (-) | - | - |
| | $\{s\}$ | -277 | - (-) | - (-) | 0.824 (0.02) | 0.883 | -6 |
| | $\{U_1, s\}$ | -218 | 0.0 (0.0) | - (-) | 0.824 (0.02) | 0.883 | -6 |
| | $\{U_2, s\}$ | -280 | - (-) | 0.221 (0.004) | 0.949 (0.007) | 0.883 | 7 |
| | $\{U_1, U_2, s\}$ | -276 | 0.0 (0.0) | 0.221 (0.004) | 0.949 (0.007) | 0.883 | 7 |
| NODDI | $\{\}$ | -204 | 0 (-) | 0 (-) | 1 (-) | - | - |
| | $\{U_1\}$ | -204 | 0.001 (0.001) | - (-) | 1 (-) | - | - |
| | $\{U_2\}$ | -247 | - (-) | 0.211 (0.006) | 1 (-) | - | - |
| | $\{U_1, U_2\}$ | -215 | 0.001 (0.001) | 0.211 (0.006) | 1 (-) | - | - |
| | $\{s\}$ | -243 | - (-) | - (-) | 0.839 (0.02) | 0.91 | -7 |
| | $\{U_1, s\}$ | -211 | 0.0 (0.0) | - (-) | 0.839 (0.02) | 0.91 | -7 |
| | $\{U_2, s\}$ | -247 | - (-) | 0.194 (0.005) | 0.945 (0.005) | 0.91 | 3 |
| | $\{U_1, U_2, s\}$ | -241 | 0.0 (0.0) | 0.194 (0.005) | 0.945 (0.005) | 0.91 | 3 |
| NODDI-DTI | $\{\}$ | -214 | 0 (-) | 0 (-) | 1 (-) | - | - |
| | $\{U_1\}$ | -214 | 0.0 (0.0) | - (-) | 1 (-) | - | - |
| | $\{U_2\}$ | -220 | - (-) | 0.085 (0.003) | 1 (-) | - | - |
| | $\{U_1, U_2\}$ | -216 | 0.0 (0.0) | 0.085 (0.003) | 1 (-) | - | - |
| | $\{s\}$ | -216 | - (-) | - (-) | 0.94 (0.017) | 0.883 | 6 |
| | $\{U_1, s\}$ | -212 | 0.0 (0.0) | - (-) | 0.94 (0.017) | 0.883 | 6 |
| | $\{U_2, s\}$ | -222 | - (-) | 0.08 (0.004) | 0.972 (0.016) | 0.883 | 10 |
| | $\{U_1, U_2, s\}$ | -213 | 0.0 (0.0) | 0.08 (0.004) | 0.972 (0.016) | 0.883 | 10 |

## References

Abdollahzadeh, A., Belevich, I., Jokitalo, E., Tohka, J., & Sierra, A. (2019). Automated 3D Axonal Morphometry of White Matter. Scientific Reports, 9, 6084. doi:10.1038/s41598-019-42648-2.

Alexander, D. C., Hubbard, P. L., Hall, M. G., Moore, E. A., Ptito, M., Parker, G. J. M., & Dyrby, T. B. (2010). Orientationally invariant indices of axon diameter and density from diffusion MRI. NeuroImage, 52, 1374–1389. URL: https://www.sciencedirect.com/science/article/pii/S1053811910007755. doi:10.1016/j.neuroimage.2010.05.043.

Beaulieu, C. (2009). CHAPTER 6 - The Biological Basis of Diffusion Anisotropy. In H. Johansen-Berg, & T. E. J. Behrens (Eds.), Diffusion MRI (pp. 105–126). San Diego: Academic Press. URL: https://www.sciencedirect.com/science/article/pii/B9780123747099000067. doi:10.1016/B978-0-12-374709-9.00006-7.

Beaulieu, C., & Allen, P. S. (1994). Determinants of anisotropic water diffusion in nerves. Magnetic Resonance in Medicine, 31, 394–400. URL: https://onlinelibrary.wiley.com/doi/10.1002/mrm.1910310408. doi:10.1002/mrm.1910310408.

Birkl, C., Langkammer, C., Haybaeck, J., Ernst, C., Stollberger, R., Fazekas, F., & Ropele, S. (2014). Temperature-induced changes of magnetic resonance relaxation times in the human brain: A postmortem study. Magnetic Resonance in Medicine, 71, 1575–1580. URL: https://onlinelibrary.wiley.com/doi/abs/10.1002/mrm.24799. doi:10.1002/mrm.24799. eprint: https://onlinelibrary.wiley.com/doi/pdf/10.1002/mrm.24799.

Burnham, K. P., Anderson, D. R., & Burnham, K. P. (2002). Model selection and multimodel inference: a practical information-theoretic approach. (2nd ed.). New York: Springer. OCLC: ocm48557578.

Campbell, J. S. W., Leppert, I. R., Narayanan, S., Boudreau, M., Duval, T., Cohen-Adad, J., Pike, G. B., & Stikov, N. (2018). Promise and pitfalls of g-ratio estimation with MRI. NeuroImage, 182, 80–96. URL: https://www.sciencedirect.com/science/article/pii/S1053811917306857. doi:10.1016/j.neuroimage.2017.08.038.

Coutu, J.-P., Chen, J. J., Rosas, H. D., & Salat, D. H. (2014). Non-Gaussian water diffusion in aging white matter. Neurobiology of Aging, 35, 1412–1421. URL: https://www.sciencedirect.com/science/article/pii/S0197458013006192. doi:10.1016/j.neurobiolaging.2013.12.001.

Cox, S. R., Ritchie, S. J., Tucker-Drob, E. M., Liewald, D. C., Hagenaars, S. P., Davies, G., Wardlaw, J. M., Gale, C. R., Bastin, M. E., & Deary, I. J. (2016). Ageing and brain white matter structure in 3,513 UK Biobank participants. Nature Communications, 7, 13629. URL: http://www.nature.com/articles/ncomms13629. doi:10.1038/ncomms13629.

Donat, C. K., Yanez Lopez, M., Sastre, M., Baxan, N., Goldfinger, M., Seeamber, R., Müller, F., Davies, P., Hellyer, P., Siegkas, P., Gentleman, S., Sharp, D. J., & Ghajari, M. (2021). From biomechanics to pathology: predicting axonal injury from patterns of strain after traumatic brain injury. Brain, 144, 70–91. URL: https://doi.org/10.1093/brain/awaa336. doi:10.1093/brain/awaa336.

Edwards, L. J., Pine, K. J., Ellerbrock, I., Weiskopf, N., & Mohammadi, S. (2017). NODDI-DTI: Estimating Neurite Orientation and Dispersion Parameters from a Diffusion Tensor in Healthy White Matter. Frontiers in Neuroscience, 11, 720. URL: https://www.frontiersin.org/article/10.3389/fnins.2017.00720. doi:10.3389/fnins.2017.00720.

Elliott, L. T., Sharp, K., Alfaro-Almagro, F., Shi, S., Miller, K. L., Douaud, G., Marchini, J., & Smith, S. M. (2018). Genome-wide association studies of brain imaging phenotypes in UK Biobank. Nature, 562, 210–216. URL: http://www.nature.com/articles/s41586-018-0571-7. doi:10.1038/s41586-018-0571-7.

Fieremans, E., Jensen, J. H., & Helpern, J. A. (2011). White matter characterization with diffusional kurtosis imaging. NeuroImage, 58, 177–188. URL: http://www.sciencedirect.com/science/article/pii/S1053811911006148. doi:10.1016/j.neuroimage.2011.06.006. Number: 1.

Frigo, M., Fick, R. H. J., Zucchelli, M., Deslauriers-Gauthier, S., & Deriche, R. (2021). Multi-Tissue Multi-Compartment Models of Diffusion MRI. bioRxiv, (p. 2021.01.29.428843). URL: https://www.biorxiv.org/content/10.1101/2021.01.29.428843v1. doi:10.1101/2021.01.29.428843. Publisher: Cold Spring Harbor Laboratory Section: New Results.

Genç, E., Fraenz, C., Schlüter, C., Friedrich, P., Hossiep, R., Voelkle, M. C., Ling, J. M., Güntürkün, O., & Jung, R. E. (2018). Diffusion markers of dendritic density and arborization in gray matter predict differences in intelligence. Nature Communications, 9, 1905. URL: https://www.nature.com/articles/s41467-018-04268-8. doi:10.1038/s41467-018-04268-8. Bandiera abtest: a Cc license type: cc by Cg type: Nature Research Journals Number: 1 Primary atype: Research Publisher: Nature Publishing Group Subject term: Human behaviour; Intelligence Subject term id: human-behaviour;intelligence.

Gong, T., Tong, Q., He, H., Sun, Y., Zhong, J., & Zhang, H. (2020). MTE-NODDI: Multi-TE NODDI for disentangling non-T2-weighted signal fractions from compartment-specific T2 relaxation times. NeuroImage, 217, 116906. URL: https://www.sciencedirect.com/science/article/pii/S105381192030392X. doi:10.1016/j.neuroimage.2020.116906.

Jelescu, I. O., Palombo, M., Bagnato, F., & Schilling, K. G. (2020). Challenges for biophysical modeling of microstructure. Journal of Neuroscience Methods, 344, 108861. URL: https://www.sciencedirect.com/science/article/pii/S0165027020302843. doi:10.1016/j.jneumeth.2020.108861.

Jelescu, I. O., Veraart, J., Fieremans, E., & Novikov, D. S. (2016). Degeneracy in model parameter estimation for multi-compartmental diffusion in neuronal tissue. NMR in Biomedicine, 29, 33–47. URL: https://onlinelibrary.wiley.com/doi/abs/10.1002/nbm.3450. doi:10.1002/nbm.3450. eprint: https://onlinelibrary.wiley.com/doi/pdf/10.1002/nbm.3450.

Jespersen, S. N., Olesen, J. L., Hansen, B., & Shemesh, N. (2018). Diffusion time dependence of microstructural parameters in fixed spinal cord. NeuroImage, 182, 329–342. URL: https://www.sciencedirect.com/science/article/pii/S1053811917306869. doi:10.1016/j.neuroimage.2017.08.039.

Jung, W., Lee, J., Shin, H.-G., Nam, Y., Zhang, H., Oh, S.-H., & Lee, J. (2018). Whole brain g-ratio mapping using myelin water imaging (MWI) and neurite orientation dispersion and density imaging (NODDI). NeuroImage, 182, 379–388. URL: https://www.sciencedirect.com/science/article/pii/S1053811917308017. doi:10.1016/j.neuroimage.2017.09.053.

Karolis, V. R., Corbetta, M., & Thiebaut de Schotten, M. (2019). The architecture of functional lateralisation and its relationship to callosal connectivity in the human brain. Nature Communications, 10, 1417. URL: https://www.nature.com/articles/s41467-019-09344-1. doi:10.1038/s41467-019-09344-1. Bandiera abtest: a Cc license type: cc by Cg type: Nature Research Journals Number: 1 Primary atype: Research Publisher: Nature Publishing Group Subject term: Cognitive neuroscience;Human behaviour;Magnetic resonance imaging;Neural circuits Subject term id: cognitive-neuroscience;human-behaviour;magnetic-resonance-imaging;neural-circuit.

Kelm, N. D., West, K. L., Carson, R. P., Gochberg, D. F., Ess, K. C., & Does, M. D. (2016). Evaluation of diffusion kurtosis imaging in ex vivo hypomyelinated mouse brains. NeuroImage, 124, 612–626. URL: https://www.sciencedirect.com/science/article/pii/S1053811915008411. doi:10.1016/j.neuroimage.2015.09.028.

Lampinen, B., Szczepankiewicz, F., Novén, M., Westen, D. v., Hansson, O., Englund, E., Mårtensson, J., Westin, C.-F., & Nilsson, M. (2019). Searching for the neurite density with diffusion MRI: Challenges for biophysical modeling. Human Brain Mapping, 40, 2529–2545. URL: https://onlinelibrary.wiley.com/doi/abs/10.1002/hbm.24542. doi:10.1002/hbm.24542. eprint: https://onlinelibrary.wiley.com/doi/pdf/10.1002/hbm.24542.

Lee, H.-H., Yaros, K., Veraart, J., Pathan, J. L., Liang, F.-X., Kim, S. G., Novikov, D. S., & Fieremans, E. (2019). Along-axon diameter variation and axonal orientation dispersion revealed with 3D electron microscopy: implications for quantifying brain white matter microstructure with histology and diffusion MRI. Brain Structure and Function, 224, 1469–1488. URL: https://doi.org/10.1007/s00429-019-01844-6. doi:10.1007/s00429-019-01844-6.

Martin Bland, J., & Altman, D. (1986). STATISTICAL METHODS FOR ASSESSING AGREEMENT BETWEEN TWO METHODS OF CLINICAL MEASUREMENT. The Lancet, 327, 307–310. URL: https://www.sciencedirect.com/science/article/pii/S0140673686908378. doi:10.1016/S0140-6736(86)90837-8.

Mohammadi, S., & Callaghan, M. F. (2021). Towards in vivo g-ratio mapping using MRI: Unifying myelin and diffusion imaging. Journal of Neuroscience Methods, 348, 108990. URL: https://linkinghub.elsevier.com/retrieve/pii/S0165027020304131. doi:10.1016/j.jneumeth.2020.108990.

Novikov, D. S., Fieremans, E., Jespersen, S. N., & Kiselev, V. G. (2019). Quantifying brain microstructure with diffusion MRI: Theory and parameter estimation. NMR in Biomedicine, 32, e3998. URL: https://onlinelibrary.wiley.com/doi/abs/10.1002/nbm.3998. doi:https://doi.org/10.1002/nbm.3998. xeprint: https://onlinelibrary.wiley.com/doi/pdf/10.1002/nbm.3998.

Novikov, D. S., Kiselev, V. G., & Jespersen, S. N. (2018a). On modeling. Magnetic Resonance in Medicine, 79, 3172–3193. URL: https://onlinelibrary.wiley.com/doi/abs/10.1002/mrm.27101. doi:10.1002/mrm.27101. eprint: https://onlinelibrary.wiley.com/doi/pdf/10.1002/mrm.27101.

Novikov, D. S., Veraart, J., Jelescu, I. O., & Fieremans, E. (018b). Rotationally-invariant mapping of scalar and orientational metrics of neuronal microstructure with diffusion MRI. NeuroImage, 174, 518–538. URL: https://www.sciencedirect.com/science/article/pii/S1053811918301915. doi:10.1016/j.neuroimage.2018.03.006.

Panagiotaki, E., Schneider, T., Siow, B., Hall, M. G., Lythgoe, M. F., & Alexander, D. C. (2012). Compartment models of the diffusion MR signal in brain white matter: A taxonomy and comparison. NeuroImage, 59, 2241–2254. URL: https://www.sciencedirect.com/science/article/pii/S1053811911011566. doi:10.1016/j.neuroimage.2011.09.081.

Sepehrband, F., Alexander, D. C., Kurniawan, N. D., Reutens, D. C., & Yang, Z. (2016). Towards higher sensitivity and stability of axon diameter estimation with diffusion-weighted MRI. NMR in Biomedicine, 29, 293–308. URL: https://analyticalsciencejournals.onlinelibrary.wiley.com/doi/abs/10.1002/nbm.3462. doi:10.1002/nbm.3462. eprint: https://analyticalsciencejournals.onlinelibrary.wiley.com/doi/pdf/10.1002/nbm.3462.

Stikov, N., Campbell, J. S. W., Stroh, T., Lavelée, M., Frey, S., Novek, J., Nuara, S., Ho, M.-K., Bedell, B. J., Dougherty, R. F., Leppert, I. R., Boudreau, M., Narayanan, S., Duval, T., Cohen-Adad, J., Picard, P.-A., Gasecka, A., Côté, D., & Pike, G. B. (2015). In vivo histology of the myelin g-ratio with magnetic resonance imaging. NeuroImage, 118, 397–405. doi:10.1016/j.neuroimage.2015.05.023.

Szafer, A., Zhong, J., & Gore, J. C. (1995). Theoretical Model for Water Diffusion in Tissues. Magnetic Resonance in Medicine, 33, 697–712. URL: https://onlinelibrary.wiley.com/doi/abs/10.1002/mrm.1910330516. doi:10.1002/mrm.1910330516. eprint: https://onlinelibrary.wiley.com/doi/pdf/10.1002/mrm.1910330516.

Takahashi, M., Hackney, D. B., Zhang, G., Wehrli, S. L., Wright, A. C., O’Brien, W. T., Uematsu, H., Wehrli, F. W., & Selzer, M. E. (2002). Magnetic resonance microimaging of intraaxonal water diffusion in live excised lamprey spinal cord. Proceedings of the National Academy of Sciences of the United States of America, 99, 16192–16196. doi:10.1073/pnas.252249999.

Tax, C. M. W., Kleban, E., Chamberland, M., Barakovíc, M., Rudrapatna, U., & Jones, D. K. (2021). Measuring compartmental T2-orientational dependence in human brain white matter using a tiltable RF coil and diffusion-T2 correlation MRI. NeuroImage, 236, 117967. URL: https://www.sciencedirect.com/science/article/pii/S1053811921002445. doi:10.1016/j.neuroimage.2021.117967.

Veraart, J., Novikov, D. S., & Fieremans, E. (2018). TE dependent Diffusion Imaging (TEdDI) distinguishes between compartmental T2 relaxation times. NeuroImage, 182, 360–369. URL: https://www.sciencedirect.com/science/article/pii/S1053811917307784. doi:10.1016/j.neuroimage.2017.09.030.

West, K. L., Kelm, N. D., Carson, R. P., Alexander, D. C., Gochberg, D. F., & Does, M. D. (2018a). Experimental studies of g-ratio MRI in ex vivo mouse brain. NeuroImage, 167, 366–371. URL: https://www.sciencedirect.com/science/article/pii/S1053811917310108. doi:10.1016/j.neuroimage.2017.11.064.

West, K. L., Kelm, N. D., Carson, R. P., Gochberg, D. F., Ess, K. C., & Does, M. D. (2018b). Myelin volume fraction imaging with MRI. NeuroImage, 182, 511–521. URL: https://www.sciencedirect.com/science/article/pii/S1053811916307935. doi:10.1016/j.neuroimage.2016.12.067.

Yu, F., Fan, Q., Tian, Q., Ngamsombat, C., Machado, N., Bireley, J. D., Russo, A. W., Nummenmaa, A., Witzel, T., Wald, L. L., Klawiter, E. C., & Huang, S. Y. (2019). Imaging G-Ratio in Multiple Sclerosis Using High-Gradient Diffusion MRI and Macromolecular Tissue Volume. American Journal of Neuroradiology, 40, 1871–1877. URL: http://www.ajnr.org/content/40/11/1871. doi:10.3174/ajnr.A6283. Publisher: American Journal of Neuroradiology Section: Adult Brain.

Zaimi, A., Wabartha, M., Herman, V., Antonsanti, P.-L., Perone, C. S., & Cohen-Adad, J. (2018). AxonDeepSeg: automatic axon and myelin segmentation from microscopy data using convolutional neural networks. Scientific Reports, 8, 3816. URL: http://www.nature.com/articles/s41598-018-22181-4. doi:10.1038/s41598-018-22181-4.

Zhang, H., Schneider, T., Wheeler-Kingshott, C. A., & Alexander, D. C. (2012). NODDI: Practical in vivo neurite orientation dispersion and density imaging of the human brain. NeuroImage, 61, 1000–1016. URL: http://www.sciencedirect.com/science/article/pii/S1053811912003539. doi:10.1016/j.neuroimage.2012.03.072. Number: 4.

